# High-order thalamic and motor cortex inputs integrate sublinearly into tuft dendrites of pyramidal neurons in mouse somatosensory cortex

**DOI:** 10.1101/2022.07.05.498894

**Authors:** Young-Eun Han, Joon Ho Choi, Kyungsoo Kim, Seongtak Kang, Hyungju Park, Ji-Woong Choi, Jong-Cheol Rah

## Abstract

Tuft dendrites of pyramidal neurons housed in layer 1 of the neocortex form extensive excitatory synaptic connections with long-range cortical and high-order thalamic axons, along with diverse inhibitory inputs. Recently, we reported that synapses from the vibrissal primary motor cortex (vM1) and posterior medial thalamic nucleus (POm) are spatially clustered together in the same set of distal dendrites, suggesting a close functional interaction. In this study, we evaluated how these two types of synapses interact with each other using in vivo two-photon Ca^2+^ imaging and electrophysiology. We observed that dendritic Ca^2+^ responses could be efficiently evoked by electrical stimulation of POm or vM1 in the overlapping set of dendritic branches, rejecting the idea of branch-wise origin-selective synaptic wiring. Surprisingly, the Ca^2+^ responses upon coincident POm and vM1 stimulation summed sublinearly. We attribute this sublinearity to mutual inhibition via inhibitory neurons because synaptic currents generated by POm and vM1 also integrated sublinearly, but pharmacologically isolated direct synaptic currents summed linearly. Inhibitory neurons receiving POm inputs in the superficial cortical layer negatively regulated vM1-evoked responses. Finally, POm and vM1 innervated overlapping but distinct populations of somatostatin-expressing inhibitory neurons. Thus, POm and vM1 inputs negatively modulate each other in the mouse somatosensory cortex.

## Introduction

The distal dendrites of pyramidal neurons in the vibrissal primary sensory cortex (vS1) are adequately positioned to integrate sensory and modulatory inputs. Despite heavy dendritic filtering, distal dendrites receive a substantial amount of excitatory synaptic inputs, including inputs from the vibrissal primary motor cortex (vM1) and the posterior medial nucleus of the thalamus (POm) (Cauller et al., 1998; Mao et al., 2011; Petreanu et al., 2009). We have demonstrated that the synapses from vM1 and POm are spatially clustered and the clusters of vM1 and POm synapses are located close to each other on the same set of dendritic branches (Kim et al., 2022), suggesting close functional interactions between the two inputs.

POm and vM1 deliver a critical set of information to vS1. The axonal boutons from vM1 in the superficial layer of vS1 showed correlated activities with various behavioral features for whisker-based object sensing, such as whisker movement and object touch (Petreanu et al., 2012). In corroboration, vM1 activity-dependent, touch-sensitive Ca^2+^ responses were observed in the distal dendrites of layer 5 (L5) pyramidal neurons (Xu et al., 2012). These findings suggest that vM1 conveys diverse behavioral information, yet the input for object touch was transmitted the best to the distal dendrites of L5 pyramidal neurons. However, information represented by POm appears to be related to whisker movement rather than object touch. POm neurons responded to facial nerve stimuli-induced “artificial active whisking” regardless of object touch (Yu et al., 2006). The responses of POm were shown to be strongly modulated in a frequency- and state-dependent manner (Diamond et al., 1992) by varying the latency and firing rate of POm neurons (Ahlssar et al., 2000). The rather complex response features of POm neurons are likely due to dense top-down inputs from vS1 and the secondary somatosensory cortex (S2) (Nothias et al., 1988; Veinante et al., 2000), besides the bottom-up inputs from the brainstem (Chiaia et al., 1991; Williams et al., 1994). Recent studies have revealed a modulatory role of POm inputs in vS1. Repetitive optogenetic POm stimulation at the frequency of near-natural whisking induced N-methyl-D-aspartate receptor (NMDAR)-dependent plateau potential in the distal dendrites, which led to rhythmic whisker deflection-dependent long-term potentiation (LTP) of sensory inputs (Gambino et al., 2014; Jouhanneau et al., 2014; Williams and Holtmaat, 2019). The potentiation of sensory inputs by POm activation appears to be mediated by intracortical inhibitory connections (Williams and Holtmaat, 2019) as well as by direct postsynaptic potentiation (Audette et al., 2019).

The inhibitory cell types that are the primary recipient of POm inputs are parvalbumin-expressing inhibitory (PV) neurons. PV neurons appear to receive the strongest and most frequent POm inputs (Audette et al., 2018; Sermet et al., 2019). Despite the sparseness of the cell population (Lee et al., 2010), vasoactive intestinal peptide-expressing (VIP) neurons and somatostatin-expressing inhibitory (SST) neurons received weak yet significant inputs in all the layers of vS1. 5-hydroxytryptamine receptor 3A-expressing inhibitory (5-HT_3a_) neurons that include VIP neurons together with neurogliaform neurons (Kawaguchi, 1995; Tamás et al., 2003) have been reported to receive stronger and more frequent POm inputs than PV neurons in the superficial layer of vS1 (Audette et al., 2018). Analogously, vM1 axons target VIP and PV neurons along with weak connections to SST neurons (Lee et al., 2013). Although the role of VIP neurons in modulating the strength of afferent sensory information has been established (Kapfer et al., 2007; Wang et al., 2004), relatively little is known about the role of SST neurons (McGarry et al., 2010). However, SST neurons are the prime candidate to modulate the branch-wise computation because they mainly target L1 and inhibit spatially constrained portions of dendritic branches (Chiu et al., 2013; Higley, 2014). Furthermore, these neurons show task-specific responses (Adler et al., 2019).

POm and vM1 inputs deliver complementary information necessary for somatosensation on the overlapping loci of the dendritic branches of pyramidal neurons and target the same inhibitory cell types. However, the integration of these two inputs into the distal dendrites of pyramidal neurons in vS1 has not been systematically explored. In the current study, we examined the integration of inputs from POm and vM1 using in vivo two-photon Ca^2+^ imaging and ex vivo whole-cell recordings to address this issue.

## Materials and Methods

### Animals

All animal experiments were conducted with the approval of the Animal Experiment Ethics Committee (approval no. IACUC-17-00031) at the Korea Brain Research Institute (KBRI). All experiments were performed using male C57BL/6J mice.

### Stereotaxic virus injection

Three-to-four-week-old male mice were anesthetized with a ketamine-xylazine mixture at 100 mg/kg ketamine-10 mg/kg xylazine and positioned in a stereotaxic apparatus (Model 942 Small Animal Stereotaxic Instrument, Kopf). Adeno-associated virus (AAV) was delivered by a glass micropipette using an automated Nanoject III injector (Drummond Scientific, Broomall, PA) following a small craniotomy. For sparse labeling of GCaMP6s, a mixture of AAV1-Syn-Flex-GCaMP6s-WPRE-SV40 and AAV1-hSyn-Cre-WPRE-hGH was co-injected at a ratio of 1:10,000 into vS1 (Anterior/Posterior (AP), −0.7; Medial/Lateral (ML), 3.5; Dorsal/Ventral (DV), 0.75 mm). For electrophysiological experiments, the mice were injected with 200 nL of AAV2-CamKIIa-hChR2(E123T/T159C)-EYFP (UNC Vector Core, Chapel Hill, NC) or 200 nL of AAV2-Syn-ChrimsonR-tdTomato (UNC Vector Core) into POm (AP, −2.06; ML, 1.25; DV, 2.97 mm) or vM1 (AP, 1.4; ML, 1.1; DV, 0.35 and 0.75 mm). For selective expression of ChR2 or fluorescent proteins in the inhibitory neurons that receive input from POm or vM1, we took advantage of the trans-synaptic anterograde viral transfer of AAV1 and the Cre-dependent double inverted open reading frame (cDIO) system. Approximately 200 nL of AAV1-hSyn-Cre-WPRE-hGH (Penn Vector Core, Philadelphia, PA) was injected into POm or vM1, and 300 nL of AAV1-mGAD67-cDIO-hChR2(H134R)-EYFP was injected into vS1 (AP, −0.7; ML, 3.5; DV, 0.35 mm). Input-dependent dual-labeling of postsynaptic neurons was achieved using the cFork anterograde tracing system (Oh et al., 2020). Then, 300 nL of AAV1-hSyn-Cre-WPRE-hGH and 300 nL of AAV1-hSyn-Flp were injected into POm and vM1, respectively, or vice versa. Subsequently, 450 nL AAV5-Ef1a-DIO-EGFP-CMV-fDIO-mScarlet was injected into vS1 (AP, −0.7; ML, 3.5; DV, 0.35 mm).

### Cranial window preparation and microelectrode implantation

Cranial window surgery was performed 21–28 days after the virus injection. Before the surgical procedure, ketoprofen (5 mg/kg) and dexamethasone disodium phosphate (5 mg/kg) were administered subcutaneously to relieve the pain and inflammation. The mice were deeply anesthetized with 1–2% isoflurane in 30% O_2_ and 70% N_2_O or ketamine/xylazine mixture. A craniotomy (approximately 3 mm diameter) was performed over the vS1 of the mouse and covered with a double-layered round glass coverslip (3 mm: 64-0720 [CS-3R], Warner Instruments, Hamden, CT; 5 mm: Electron Microscopy Sciences, Hatfield, PA, 72195-05), bonded with an optical adhesive (NOA71, Norland, Jamesburg, NJ). Stimulation electrodes (concentric electrodes, TM33CCINS, World Precision Instruments, Sarasota, FL) were implanted in POm and vM1 through small holes (approximately 0.7 mm) in the skull at 35° and 45° angles relative to the vertical axis, respectively. A customized head plate was attached to the skull with dental cement (Super-Bond C&B kits, Sun Medical, Shiga, Japan). The mice had at least 4 h for recovering from surgery before in vivo two-photon Ca^2+^ imaging.

### In vivo two-photon calcium imaging and detection

Two-photon imaging was performed using a Ti:sapphire laser (Chameleon, Coherent, Santa Clara, CA) equipped with an x-y galvanometer scanning system (Scientifica, East Sussex, UK). GCaMP6s calcium indicator was excited at 910 nm (typically 20–40 mW power under the objective for apical tuft imaging) and imaged through Olympus 40x water immersion lens (Olympus, Shinjuku, Japan, 0.80 N.A., 3.3 mm working distance). Emission light was passed through a 565 DCXR dichroic filter (Chroma Technology, Bellows Falls, VT) and an ET525/70m-2p filter (Chroma Technology) and was detected using a GaAsP photomultiplier tube (MDU-PMT-50-50, Scientifica). Images with 256 × 256 pixels were acquired at 4 Hz using Labview-based image acquisition software SciScan (Scientifica).

Each imaging trial consisted of 20 or 30 frames. In each trial, three 1 ms-long stimuli were delivered (50 Hz) at the 5^th^ frame through concentric bipolar microelectrodes (FHC, Bowdoin, ME) in POm, vM1, or both. We repeated 20 or 30 trials for each imaging condition. Event-related regions of interest (ROI) were detected from the motion-corrected two-photon image series using custom-built MATLAB scripts (Mathworks, Natick, MA). Specifically, to minimize motion-caused artifacts, we aligned image frames using a non-rigid motion correction algorithm (Pnevmatikakis and Giovannucci, 2017). We detected clusters of pixels that showed expected fluorescence changes in response to the stimulus. In each trial, the pixel intensity difference (PF) was normalized to the baseline acquired from the average Ca^2+^ intensity before stimulation onset (PF_0_) as (PF – PF_0_)/PF_0_. Then, the stimulus-modulated pixels were defined as those with a statistically significant projection of fluorescence deflection into the template modeled by a double exponential function (*p* < 0.05, one-tailed Student’s t-test). The intensities of the clustered pixels were averaged to estimate the fluorescence in the ROIs. The detected ROI traces were denoised and used for ΔF/F_0_ estimation, which was calculated as (F – F_0_)/F_0_, where F is the ROI trace in a trial and F_0_ is the ROI baseline, which is the mean intensity of the pixels in the frames before stimulation onset.

### Brain slice preparation

Acute brain slices were prepared from 7-9-weeks old male C57BL/6J mice. After anesthetizing the mice with sodium pentobarbital at 70 mg/kg, transcardial perfusion was performed with 25 mL of approximately 25 °C cutting solution containing the following (in mM): 93 NMDG, 2.5 KCl, 1.2 NaH_2_PO_4_, 30 NaHCO_3_, 20 HEPES, 20 glucose, 2 thiourea, 5 sodium ascorbate, 3 sodium pyruvate, 0.5 CaCl_2_, 10 MgCl_2_, and 12 N-acetyl-L-cysteine with pH adjusted to 7.4 by adding HCl. The mice were decapitated, and their brains were quickly removed and chilled in the cutting solution. The isolated brain was glued onto a vibratome (VT1200S, Leica, Wetzlar, Germany) stage angled at 10°, and 300 μm-thick coronal brain slices were acquired. The slices were incubated at 32 °C for 15 min in the recovery solution containing the following (in mM):104 NaCl, 2.5 KCl, 1.2 NaH_2_PO_4_, 24 NaHCO_3_, 10 HEPES, 15 glucose, 2 thiourea, 5 sodium ascorbate, 3 sodium pyruvate, 2 CaCl_2_, 2 MgCl_2_, and 12 N-acetyl-L-cysteine and thereafter maintained at approximately 25 °C in the artificial cerebrospinal fluid (aCSF) containing the following (in mM): 125 NaCl, 2.5 KCl, 1.2 NaH_2_PO_4_, 24 NaHCO_3_, 5 HEPES, 13 glucose, 0.4 sodium ascorbate, 2 sodium pyruvate, 2 CaCl_2_, and 1 MgCl_2_. All solutions were saturated with carbogen (95% O_2_ and 5% CO_2_) to a final pH of 7.4.

### In vitro electrophysiology and optogenetic stimulation

L5 pyramidal neurons of vS1 were visualized using an upright microscope equipped with differential interference contrast optics (BX51WI, Olympus) with a 40x water immersion objective (NA 0.8, Olympus). All electrophysiological recordings were performed at 30 °C, and fresh aCSF was perfused at approximately 1.5 mL/min. The patch electrodes were pulled from borosilicate glass capillaries to obtain a resistance between 3 and 4 MΩ. The internal solution contained the following (in mM):138 potassium gluconate, 10 KCl, 10 HEPES, 10 Na_2_-phosphocreatine, 4 MgATP, 0.3 NaGTP, and 0.2 EGTA (pH 7.25). Electrophysiological recordings from soma were generated using a MultiClamp 700B amplifier and Digidata 1550 (Molecular Devices, San Jose, CA) at a sampling rate of 10 kHz and acquired using a PClamp 10 (Molecular Devices). EPSCs and IPSCs were recorded at a holding potential of −60 to −70 mV and 0 mV, respectively. EPSCs and IPSCs were pharmacologically verified. EPSCs were completely abolished in the presence of 25 μM AP5 (Tocris, Bristol, UK) and 50 μM CNQX (Tocris), while IPSCs were completely inhibited by the GABA_A_ receptor antagonist bicuculline (Sigma-Aldrich, St. Louis, MO). To assess synaptic input from either or both POm and vM1, optic fibers were placed at L1 of vS1, where apical tuft dendrites of the recorded L5 pyramidal neurons were located. POm, vM1, or both axons expressing ChR2 or ChrimsonR were stimulated optogenetically with 1–20 ms light pulses from either or both 473 nm (BL473T3-100FC-ADR-800A, SLOC) and 589 nm DPSS laser (MGL-F-589, CNI). The data were analyzed using Clampfit 10.4 (Molecular Devices) and IGOR Pro software (Wavemetrics, Portland, OR).

### Quantification of the linearity

The linearity index was defined as follows:

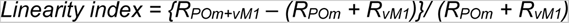

where R_POm+vM1_ stands for the size of the responses, such as Ca^2+^ responses or EPSCs evoked by simultaneous POm and vM1 stimulation, and R_POm_ and R_vM1_ are the sizes of the responses evoked by POm or vM1 stimulation, respectively. To compare the response by simultaneous stimulation of POm and vM1 with the larger response of either of the two responses (Max), the multi-input enhancement (ME) index was devised as follows:

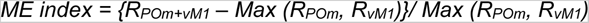

Max (a, b) is defined as the greater response between a and b, and R represents the size of the response to the subscribed stimulation.

### Immunohistochemistry

Mice were deeply anesthetized with sodium pentobarbital at 70 mg/kg and perfused transcardially with phosphate-buffered saline followed by 4% paraformaldehyde (PFA). Brains were removed and fixed with 4% PFA overnight at 4 °C. The brain was sectioned into 50 μm-thick slices using a vibratome (VT1200S, Leica). After washing in Tris-buffered saline (TBS), brain slices were permeabilized with 0.3% Triton X-100 (Fisher scientific, Waltham, MA, BP151-500) in TBS (TBS-T) for 30 min, followed by three washes in TBS, and transferred into blocking buffer, 5% donkey serum (Jackson ImmunoResearch, West Grove, PA, 017-000-121) in TBS-T for 1 h at approximately 25 °C. The sections were then incubated with primary antibodies against parvalbumin (1:500, Synaptic system, Goettingen, Germany, 195 004), somatostatin (1:50, Santa Cruz Biotechnology, Dallas, TX, sc-47706), vasoactive intestinal peptide (VIP) (1:100, Immunostar, 20077), or GFP (1:500, Abcam, Cambridge, UK, ab1218) overnight at 4 °C and rinsed three times with 0.3% tween-20 (P9416, Sigma-Aldrich) in TBS (TBS-T). The brain slices were subsequently incubated in blocking buffer (5% donkey serum in TBS-T) containing appropriate fluorescence-conjugated secondary antibodies (1:1000; Jackson ImmunoResearch; Dylight 405-conjugated donkey anti-guinea pig IgG [706-475-148], Alexa Fluor 647-conjugated donkey anti-rat IgG [712-605-150], Alexa Fluor 594-conjugated donkey anti-rabbit IgG [711-585-152], and Alexa Fluor 488-conjugated donkey anti-mouse IgG [715-545-150]) for 1 h at approximately 25 °C. After washing five times in TBS, the slices were mounted using 200 μL Prolong diamond antifade mounting solution (Invitrogen, Waltham, MA, P36961) on micro slide glasses (Matsunami glass, Osaka Prefecture, Japan, S959502). Images were acquired using a confocal microscope (TI-RCP, Nikon, Tokyo, Japan).

### Statistical analysis

Statistical tests were performed using OriginPro 2018 (OriginLab, Northampton, MA), GraphPad Prism 8.4.3 (GraphPad), or MATLAB (MathWorks). Data are expressed as mean ± S.E.M., and statistical differences between the experimental groups were analyzed using paired two-sample Student’s t-test. A *p*-value < 0.05 was considered statistically significant. The Kolmogorov–Smirnov test was performed to determine the statistical differences in the cumulative distributions.

## Results

### Identification of the dendrites receiving inputs from vM1 and POm

To examine the functional interactions between the inputs from vM1 and POm in the distal dendrites of L5 pyramidal neurons of vS1, we transduced the L5 pyramidal neurons with GCaMP6s using adeno-associated virus (AAV) (Figure 1A). To identify the dendritic patches that show Ca^2+^ responses by both vM1 and POm stimulation, we measured Ca^2+^ deflection in the dendritic branches in layer 1 (L1) (at approximately 10 μm from the cortical surface) with electrical stimulation of vM1 or POm. At the 5^th^ frame of the 20-frame scanning, three consecutive electrical stimuli were delivered at 50 Hz to boost the chance of successful synaptic transmission (Figure 1B, see the Materials and Methods section for further details). We identified dendrites that showed rapid and reliable Ca^2+^ deflection upon the stimulation (Figure 1D-E and Supplementary Figure 1). The size of the responses varied, but responses evoked by vM1 stimulation usually resulted in a greater Ca^2+^ amplitude and success rate than those evoked by POm stimulation (Figure 1F-H). This observation corroborates our anatomical finding that the number of vM1 synapses is around twice the number of POm synapses in the distal dendrites (Kim et al., 2022). However, responses by POm stimulation lasted longer than those by vM1 (Figure 1I). Because of the different response kinetics between the inputs, we defined the strength of the responses as the total area of the Ca^2+^ responses instead of the peak amplitude.

**Figure 1.**
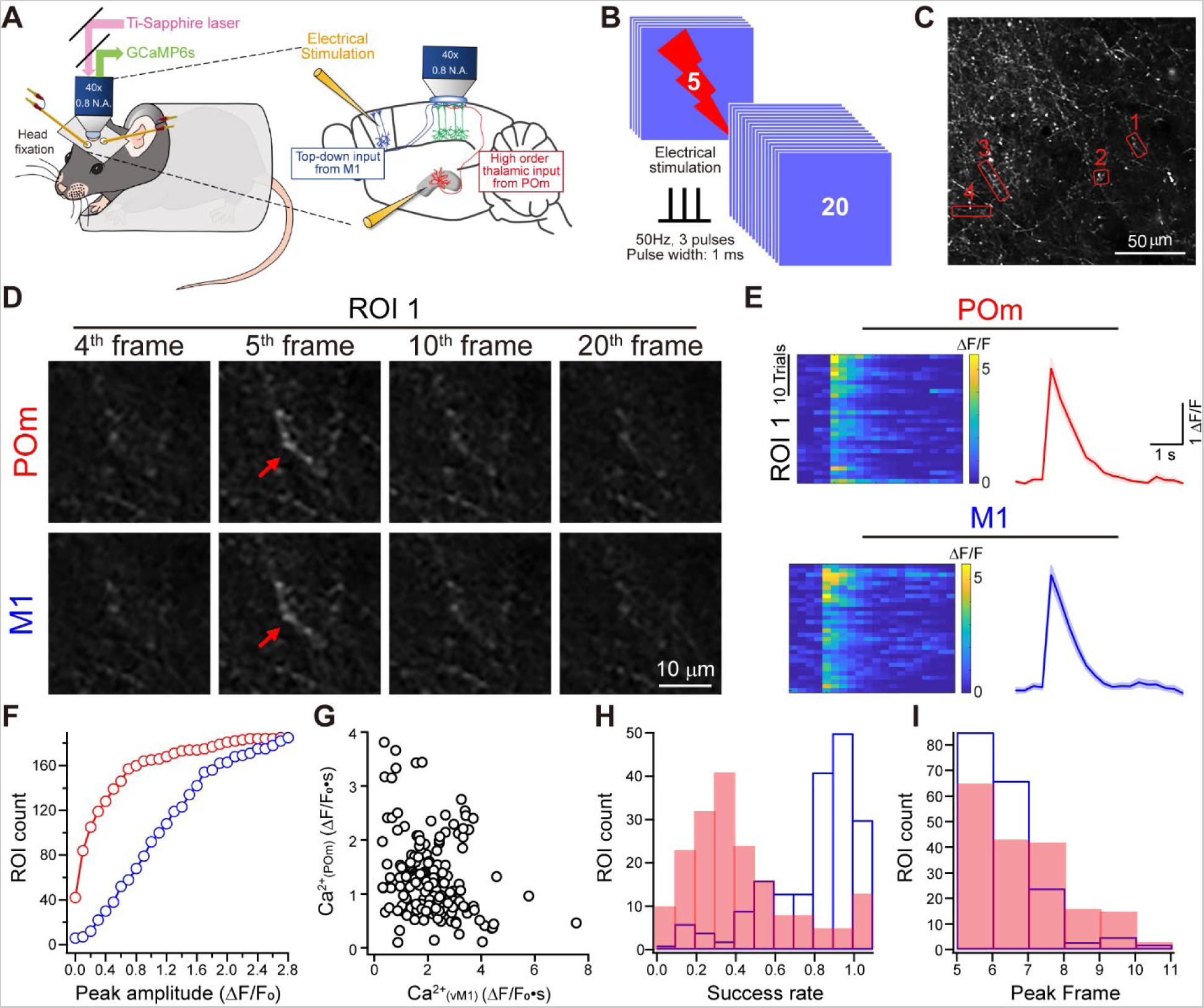
Identification of the dendritic branches responding to both vM1 and POm stimulation. **A-B.** Experimental schematic. GCaMP6s was expressed sparsely in L5 of vS1, and stimulation electrodes were implanted in POm and vM1 of the mice. Distal dendritic Ca^2+^ responses upon electrical POm or vM1 stimulation were imaged through a cranial window (A). Twenty frames were scanned at 4 Hz per trial and a set of electrical stimuli was delivered at the 5^th^ frame (B). **C.** Clusters of pixels that respond to the electrical stimulations in synchrony were selected as ROIs, and ΔF/F_0_ was calculated for each ROI. ROIs with expected ΔF/F_0_ change were shortlisted (red boxes with numbers). **D**. Representative images of baseline (4^th^), immediately after stimulation (5^th^), and the following responses (at 10^th^ and 20^th^ frames), from a dendritic branch responding to POm (upper) and vM1 (lower) stimulation. Images are from the red box 1 in C. See the Supplementary Figure 3 for ROIs 2–4. **E**. Color-coded ΔF/F_0_ for 30 repetitive trials (left) and averaged ΔF/F. Shaded areas represent standard errors of the response. **F**. Cumulative histogram of trial averaged peak ΔF/F_0_ of ROIs. **G**. The relative response strength upon the stimulation of POm and vM1 for all ROIs (n = 185). **H**. Histogram of the success rates in ROIs. **I.** Histogram of the frames that showed the peak ΔF/F_0_.

### Linearity of the distal Ca^2+^ responses by vM1 and POm stimulation

Having identified the dendritic patches that respond to both vM1 and POm stimulation, we examined the functional interactions between the two inputs. We measured Ca^2+^ deflection by simultaneous stimulation of vM1 and POm and compared it with the arithmetic sum of the responses by individual stimulation (Figure 2). In many branches, we observed that simultaneous stimuli evoked significantly smaller Ca^2+^ responses than expected arithmetically (Figure 2). Surprisingly, some dendritic branches showed an even smaller response to simultaneous stimuli than to a single stimulus (Figure 2A-C). To quantify the linearity of the integration, we devised a parameter called *linearity index*, defined as the difference of the response from arithmetic expectation normalized by the sum of the two responses by individual stimulation (Materials and Methods). The linearity index is zero if the two inputs are integrated linearly, whereas super- and sub-linear integrations show positive and negative values, respectively. On average, the dendritic patches showed a negative linearity index (−0.27 ± 0.03, median = −0.325, Figure 2D). The saturation of Ca^2+^ could not explain this phenomenon in the dendritic patches for the following reasons. The multi-input enhancement (ME) index, defined as Ca^2+^ responses to the simultaneous stimuli normalized by the larger response of single input, was near to but less than zero (0.09 ± 0.04, median = −0.025). This suggests that simultaneous stimuli evoked a smaller response than single-site stimuli. Sublinearity was observed regardless of the input strength (Figure 2C) or the strength difference between the two inputs (Figure 2E). We classified the dendritic patches into three groups, depending on the relative strength of the two inputs in each dendritic patch, using hierarchical clustering analysis (Supplementary Figure 2). With the center defined along the line where the vM1 response was twice as strong as the POm response, the responses were classified above and below the line (Supplementary Figure 2A-B). We found that sublinearity occurred regardless of the relative strength difference, supporting the conclusion drawn from the linearity index (Supplementary Figure 2C-D). Thus, we concluded that the inputs from vM1 and POm are integrated sublinearly in the distal dendrites of pyramidal neurons, independent of the input strength or the strength difference between the two inputs.

**Figure 2.**
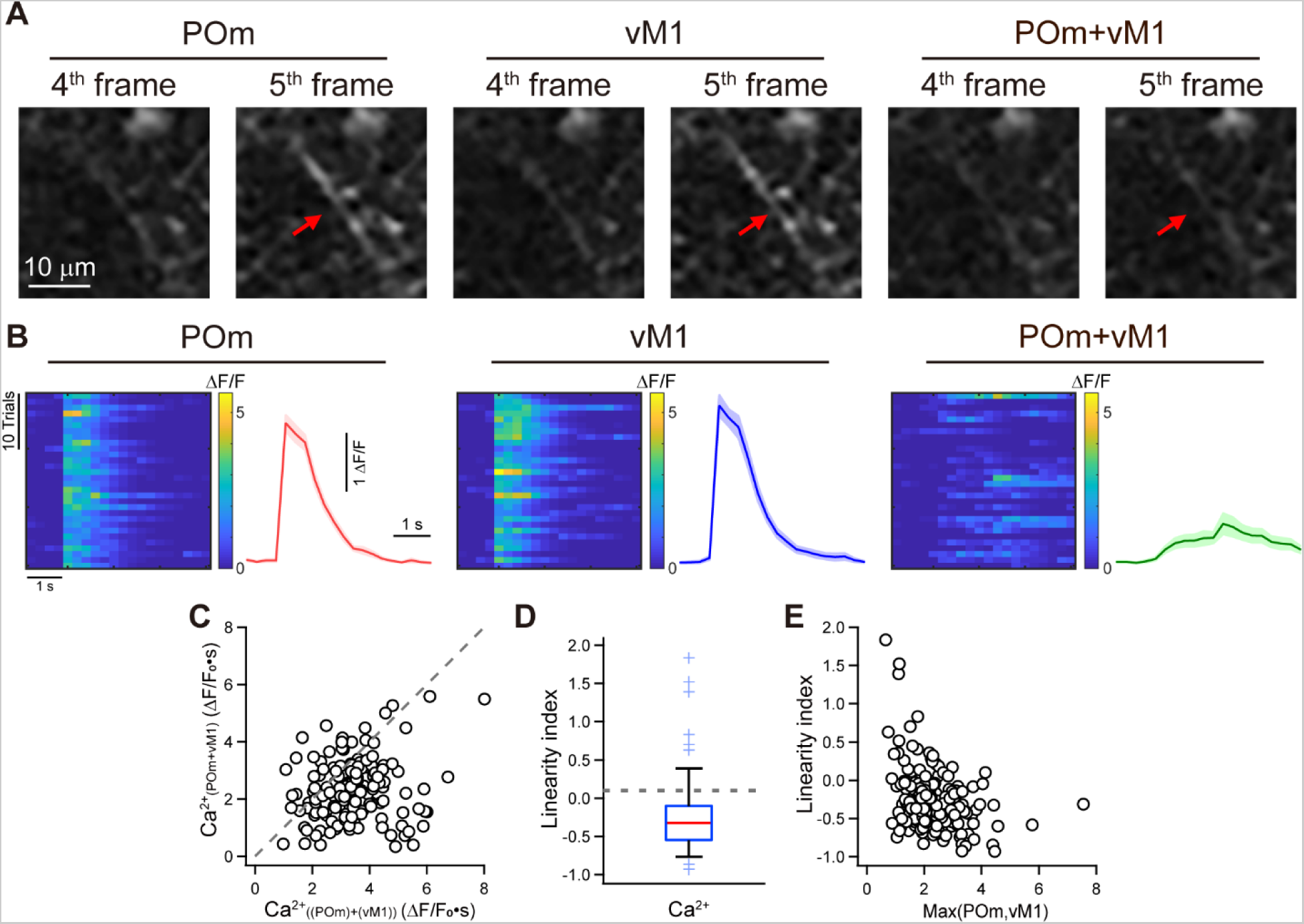
Summation of the calcium deflection by the stimulation of POm and vM1. **A.** Representative dendrites with strong sublinear input integration. Frame images immediately before and after stimulation of POm (left) and vM1 (middle) and simultaneous POm and vM1 stimulation (right) at a representative ROI are shown in Figure 1C. **B**. Ca^2+^ responses of the exemplified dendrites in A. ΔF/F_0_ for 30 repetitive trials and averaged ΔF/F_0_ by stimulation of POm (left), vM1 (middle), and both (right) from the representative ROI. **C**. Population Ca^2+^ deflection by simultaneous POm and vM1 stimulation as a function of the linear expected value by individual stimulations (n = 185). **D**. Linearity index of response integration, defined as {ΔCa_POm+vM1_ − (ΔCa_POm_ + ΔCa_vM1_)}/ (ΔCa_POm_ + ΔCa_vM1_). **E**. Linearity index as a function of strength of the larger Ca^2+^ responses. Scale bars = 200 µm; scale bars in insets= 10 µm.

### Summation of the synaptic inputs from vM1 and POm using whole-cell recording

Having established the sublinearity of the Ca^2+^ response integration, we wondered whether sublinear integration could be observed at the level of synaptic transmission. We delivered channelrhodopsin-2 (ChR2) and ChrimsonR to POm and vM1, respectively, or vice versa, via an AAV (Figure 3A). We then measured the optogenetically evoked excitatory postsynaptic currents (oEPSCs) by selectively illuminating the top layer of vS1 (Figure 3A and Supplementary Figure 3). Similar to the integration of the dendritic Ca^2+^ responses, we found that the oEPSC evoked by simultaneous vM1 and POm excitation (oEPSC_vM1+POm_) was significantly smaller than the arithmetic sum of the oEPSC evoked by POm stimulation (oEPSC_POm_) and oEPSC evoked by vM1 stimulation (oEPSC_vM1_) (Figure 3B and C, oEPSC_vM1+POm_ = 158.74 ± 41.65 pA, arithmetic sum = 197.30 ± 49.36 pA, *p* = 0.0029, n = 14, paired t-test). The linearity index of the synaptic response showed a negative value (−0.21 ± 0.03, median = −0.178, Figure 3H). As observed for the dendritic Ca^2+^ responses, sublinearity was independent of the input strength (Figure 3F) or the difference between the strength of the inputs from the two regions (Figure 3G). This finding suggests that the sublinear integration of the oEPSCs occurs at the level of distal synaptic transmission, at least in part.

**Figure 3.**
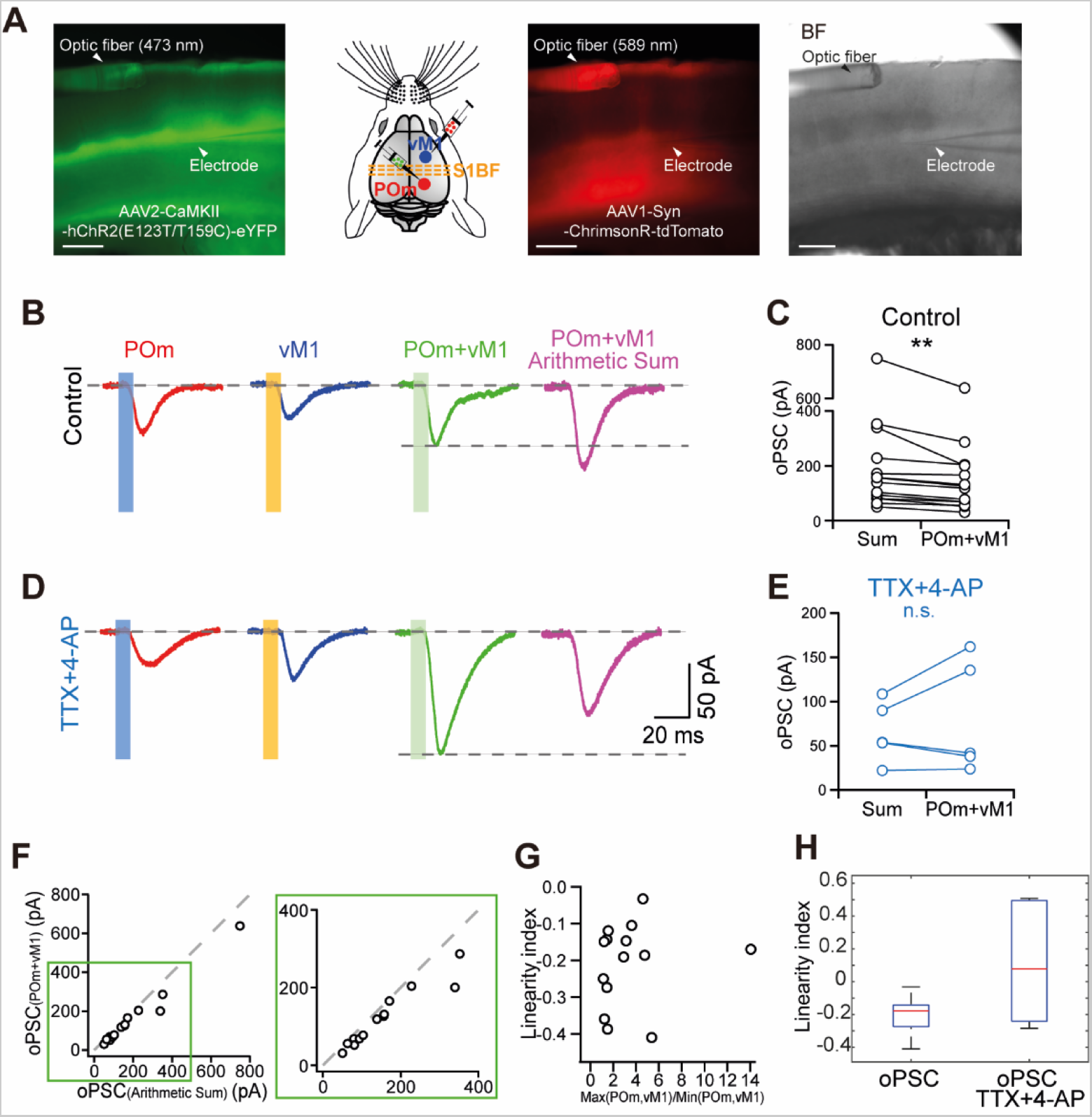
Integration of the synaptic current evoked by POm and vM1. **A.** Experimental setup. Adeno-associated virus (AAV)-transducing hChR2-eYFP and ChrimsonR-tdTomato was stereotactically delivered to POm and vM1, respectively, or vice versa. Excitation lights (473 and 589 nm) were guided toward the top layer of vS1 via an optic fiber, while postsynaptic currents were recorded from the soma of the L5 pyramidal neurons of vS1. From the left, POm inputs expressing hChR2-eYFP, a schematic of viral delivery for optogenetic stimulation of the inputs, ChrimsonR-tdTomato expressed in vM1, and a bright-field image of the experimental setup are shown. **B.** Example traces of oPSCs generated by the stimulation of POm (red), vM1 (blue), or both (green) and the arithmetic sum of individually evoked POm and vM1 oPSCs (magenta). **C.** Population comparison between the sum of individually evoked oPSCs and oPSC by simultaneous stimulation. **D-E.** Similar to B-C, except that oEPSCs were measured in the presence of 1 µM tetrodotoxin (TTX) and 100 µM 4-aminopyridine (4-AP). **F.** oPSCs generated by simultaneous POm and vM1 stimulation compared to the sum of oPSCs generated by individual POm and vM1 stimulations. (Inset) Zoomed at 0–400 pA. **G.** Relationship between the linearity index and ratio of the two inputs. **H**. Linearity indices with and without polysynaptic contributions. Scale bar = 200 μm

### Summation of the synaptic inputs without feedforward inhibition

A possible underpinning of the significant degree of sublinear summation is the involvement of inhibitory neurons. Specifically, we tested whether the excitatory inputs from POm may inhibit the transmission of the vM1 inputs through feedforward inhibition and vice versa. Such mutual inhibition between the inputs could account for the suppressed Ca^2+^ responses by simultaneous stimulation, as exemplified in Figure 2B. To test this possibility, we first examined the existence of feedforward inhibition at the level of distal dendrites of L5 pyramidal neurons. To measure the distal inputs from vM1 and POm selectively, we delivered the excitation light on the surface layer of vS1 in brain slices (Supplementary Figure 3). In concordance with previous observations (Audette et al., 2018; Lee et al., 2013), we found that optogenetic stimulation of distal inputs from vM1 and POm evoked excitatory as well as inhibitory synaptic currents on the pyramidal neurons of vS1 (Supplementary Figure 4). The onset of optogenetically evoked inhibitory postsynaptic currents (oIPSCs) was significantly slower than that of oEPSCs (Figures 4B and D), which suggested feedforward inhibition. The onset of oEPSCs measured at the reversal potential of chloride (Cl^−^) was approximately 5 ms after the stimulation (5.11 ± 1.24 ms for POm and 5.21 ± 0.35 ms for vM1), while the onset of oIPSCs measured at 0 mV was further delayed (7.34 ± 1.46 ms for POm and 8.25 ± 0.86 ms for vM1, *p* = 0.0064 for POm, 0.0018 for vM1, paired t-test). Occasionally, at 0 mV, the remaining inward oEPSCs were observed, and temporally delayed inhibitory inputs were clearly visualized (Figure 4C). Consequently, to directly examine whether feedforward inhibition is the source of the sublinear summation, we measured the linearity of the integration without the involvement of inputs from polysynaptically excited neurons. To isolate the action potential (AP)-independent monosynaptic transmission, we blocked Na^+^ channels with 1 μM tetrodotoxin (TTX) and delayed repolarization with 100 μM 4-aminopyridine (4-AP) to allow optogenetic depolarization to evoke neurotransmitter release (Petreanu et al., 2009). We found that the oEPSC was linearly integrated in this condition (80.24 ± 28.39 pA for oEPSC_POm+vM1_ vs. 65.61 ± 15.17 pA for arithmetic sum, *p* =0.37, n = 5, paired t-test, Figure 3D, E, and H). This finding demonstrated that the sublinearity of the oEPSC is mediated by polysynaptic inhibition, at least in part, and suggested that feedforward inhibition may function as mutual inhibition.

**Figure 4.**
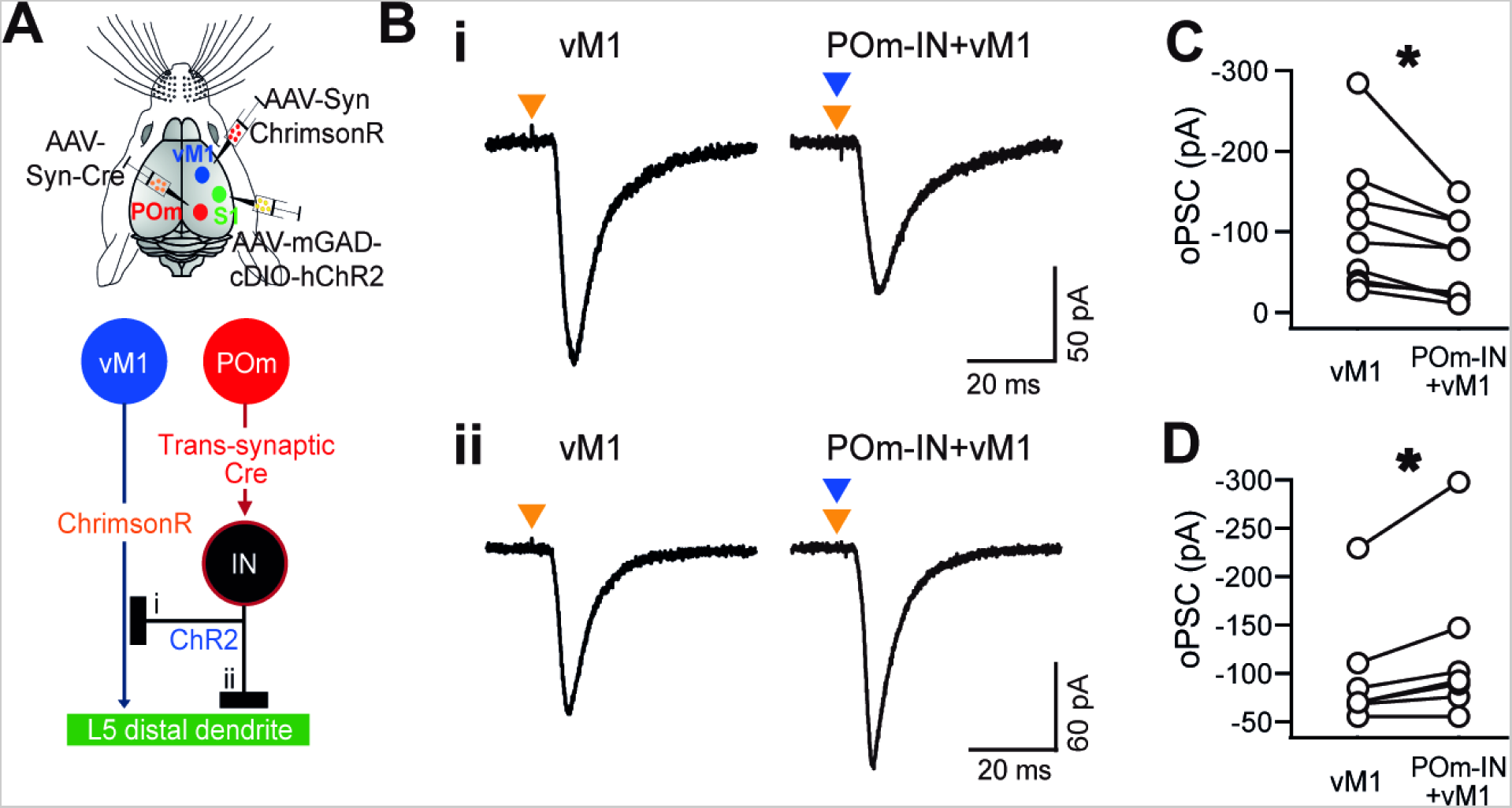
Identification of the inhibition targets. **A.** Schematic representation of the experiment. Trans-synaptic AAV1-hSyn-Cre-WPRE-hGH was injected into POm, whereas AAV1-mGAD67-cDIO-hChR2(H134R)-EYFP was injected into vS1 for selective ChR2 expression in the inhibitory neurons receiving POm inputs (POm-IN) in vS1. AAV2-Syn-ChrimsonR-tdTomato was delivered to vM1. As shown in Figure 3, excitation light (473 and 589 nm) was guided toward the top layer, while postsynaptic currents were recorded at the membrane potential, where the excitation of POm-IN generated an inward current. The two possible targets of inhibition are (i) presynaptic vM1 axons or (ii) postsynaptic structures. **B**. Example traces of scenarios (i) and (ii) by the excitation of vM1 with and without simultaneous excitation of POm-IN. **C-D**. Comparison of oPSC_vM1_ populations in the absence and presence of POm-IN stimulation. Note that simultaneous POm-IN stimulation with an internal solution allowing inward Cl^−^ current decreases oPSC_vM1_ in 10 out of 18 neurons (C) and increases it in the rest (D).

### Modulation of vM1 inputs by the POm-driven feedforward inhibition

Our data indicated that feedforward inhibition is involved in sublinear integration. However, the target of the inhibition was unclear. While we measured the inhibitory inputs directly from the postsynaptic neurons (Supplementary Figure 4), we observed sublinear synaptic transmission near the equilibrium potential of Cl^−^ (Figure 3), although the degree of sublinearity was more moderate, indicating that inhibition had to occur presynaptically. To address this, we expressed ChrimsonR in vM1 and delivered AAV1-Cre to POm, while mGAD67-cDIO-ChR2-EYFP was expressed in vS1. Thus, trans-synaptically transferred Cre recombinase from POm will be expressed only in the inhibitory neurons. We then voltage-clamped the pyramidal neurons with an internal solution in which chloride conductance generated an excitatory current. In this experimental setup, presynaptically targeted inhibitory inputs from the inhibitory neurons receiving POm inputs or POm-IN (oIPSC_POm-IN_) decreased the oEPSC_vM1_ by reducing the depolarization (Figure 4Ai). In contrast, postsynaptically targeted oIPSC_POm-IN_ added the excitatory current on top of the EPSC_vM1_ (Figure 4Aii). At approximately −65 mV (−65.38 ± 1.19 mV), optogenetic stimulation of POm-IN (oIPSC_POm-IN_) generated a bicuculline-dependent inward current (−19.46 ± 7.66 pA). Upon the simultaneous stimulation of POm-IN and vM1 inputs, we observed decreased synaptic current in 9 out of 16 cells (EPSC_vM1_ = −104.93 ± 27.69 pA; EPSC_vM1_ with IPSC_POm-IN_ = −67.32 ± 17.13 pA, *p* = 0.02, paired t-test, n = 9), whilst increased synaptic currents were observed in the rest (EPSC_vM1_ = −98.18 ± 22.83 pA vs. EPSC_vM1_ with IPSC_POm-IN_ = −122.83 ± 31.03 pA, *p* = 0.03, paired t-test, n = 7). We confirmed that the changes were bicuculline dependent in all the recorded neurons (Supplementary Figure 5). We concluded that the excitatory vM1 inputs are indeed modulated by the inhibitory neurons that receive POm inputs and that the feedforward inhibition targets predominantly postsynaptic neurons but can also target presynaptic structures.

### Interneuron population mediating the feedforward inhibition

So far, we have described a substantial sublinear summation of vM1 and POm inputs in vivo and in vitro. The sublinearity was partially mediated by feedforward inhibition. However, for such wiring to exert mutual inhibition, POm and vM1 axons must target a different population of inhibitory neurons. To test this hypothesis, we first examined whether vM1 and POm axons target different types of inhibitory neurons. We identified the inhibitory neurons receiving inputs from vM1 or POm by the anterograde trans-synaptic delivery of AAV1-Cre and Cre-dependent ChR2 with enhanced yellow fluorescent protein (EYFP) expression under the control of the *Vgat* promoter in vS1 (Zingg et al., 2017). We then identified the cell types of the inhibitory neurons by immunohistochemistry to detect PV, SST, and VIP neurons (Figures 5A and C). We identified SST (42.15 ± 2.07 % for POm, 62.61 ± 3.13 % for vM1) and PV-expressing (22.81 ± 1.58 % for POm, 19.07 ± 2.24 % for vM1) neurons among the POm- or vM1-connected GABAergic neurons. However, VIP neurons were found scarcely (0.97 ± 0.59 % for POm, 5.25 ± 0.87 % for vM1, Figure 5A-D). As both inputs target the overlapping population of inhibitory neurons, we rejected the possibility that the two inputs may target distinct inhibitory cell types. We then examined whether the populations of SST or PV neurons that received inputs from vM1 and POm were distinct. To identify the population of neurons that received the inputs, we performed AAV1-mediated anterograde trans-synaptic delivery of Cre or Flip and dual gene expression cassette under the control of the *Vgat* promoter in vS1 (Oh et al., 2020). Inhibitory neurons receiving POm inputs expressed EGFP, whereas those connected with vM1 expressed mScarlet (Figure 5E-G). Both PV and SST neurons that received the vM1 and POm inputs were spread over all the layers (Figure 5A and F). We noticed that some pyramidal neurons also expressed EGFP, mScarlet, or both, probably because of leaky control of the *Vgat* promoter.

**Figure 5.**
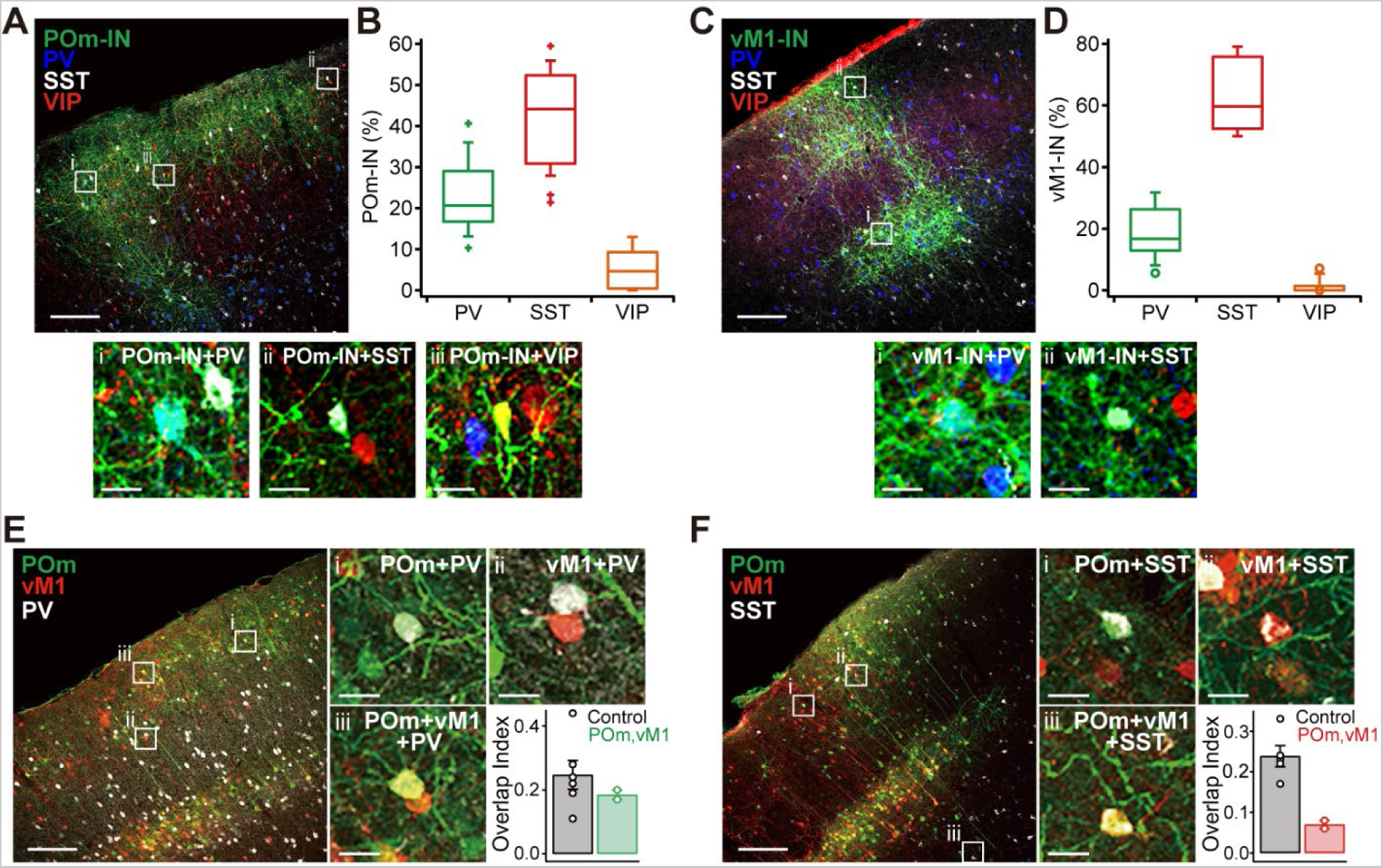
The population of inhibitory neurons that mediate feedforward inhibition. **A-D.** The identification of inhibitory neurons receiving inputs from POm, vM1, or both. **A.** Maximum intensity projected confocal images of vS1. POm-connected neurons labeled by trans-synaptic labeling (green) and immunostained PV (blue), SST, and VIP neurons. **B**. Proportion of cell types out of POm-connected, PV, SST or VIP neurons. POm-connected neurons without co-labeling of the PV, SST, or VIP neurons were excluded from the calculation. **C**. Similar to A, the vM1-connected neurons are labeled in green. **D**. Cell type population of vM1-connected inhibitory neurons. **E-F.** Fraction of PV (E) or SST (F) neurons that received both POm and vM1 inputs. **E.** Maximum intensity projected confocal image of vS1 with trans-synaptically labeled POm-connected (green) and vM1-connected (red) inhibitory neurons, and immunostained PV neurons (white). The zoomed-in view of POm-connected (i) and vM1-connected (ii) PV neurons, and PV neurons receiving both the inputs (iii). The fraction of PV neurons receiving both inputs to the PV neurons receiveing either input (overlap index) or iii/(i+ii+iii). **F.** Same as E, but SST neurons are shown in white color. Scale bars = 200 and 20 μm.

However, none of the neurons with pyramidal cell morphology were immunostained for PV or SST (Figure 5E and F). We devised an overlap index parameter defined as the fraction of the neurons expressing both EGFP and mScarlet out of the interneurons expressing either of the fluorescent proteins. The overlap of the inputs tends to be underestimated owing to the false-negative labeling of AAV1-mediated neurotracers; the neurons receive synaptic inputs but have no neurotracer label. To control for this, we compared the overlap index of the two inputs with the overlap index when the two recombinases were transduced at the same site (vM1 or POm, Figure 5E and F control). We found that the overlap index of SST neurons was appreciably lower than that of the control (0.24 ± 0.03 for same site control vs. 0.07 ± 0.01 for SST), suggesting SST neurons preferentially received inputs from POm or vM1 inputs. The low overlap index cannot be explained by the false-negative labeling of the neurotracers alone because the overlap index for PV neurons was not significantly different from that of the same site control (0.23 ± 0.02 for the same site control and 0.18 ± 0.01 for POm and vM1). Our histological analyses indicated that vM1 and POm target overlapping inhibitory cell types. However, the populations of SST neurons targeted by vM1 and POm are segregated more randomly in vS1.

Taken together, we demonstrated a mode of sublinear synaptic integration in the distal dendrites of vS1 through feedforward inhibition and suggested that the inhibitory inputs from POm and vM1 can be attributed to the segregated population of SST neurons.

## Discussion

We explored the underlying branch-specific dendritic responses observed in behaving animals (Cichon and Gan, 2015; Xu et al., 2012). In our previous study, we assessed the possibility of branch-selective wiring of inputs on distal dendrites. To reject the possibility, we found that the synapses from POm and vM1 were highly intertwined (Kim et al., 2022). In the current study, we explored the functional interactions between the two inputs to better understand how the intermingled synaptic populations of different origins may reconcile with the reported branch-specific dendritic responses (Cichon and Gan, 2015; Xu et al., 2012). We first re-evaluated the pattern of synaptic wiring within dendritic branches using an independent experimental method. Excitation of Pom- or vM1-evoked Ca^2+^ responses in a largely overlapping population of dendrites and the responses evoked by vM1 were approximately two-fold larger than those by POm, corroborating with our anatomical observation that the number of vM1 synapses were twice higher than that of POm synapses (Figure 1). The time-to-peak of the Ca^2+^ responses by POm excitation was slower than that by vM1 excitation, likely due to the strong NMDAR-dependent component of the synaptic potential in POm synapses (Gambino et al., 2014; Jouhanneau et al., 2014; Williams and Holtmaat, 2019). We found that the inputs from POm and vM1 were integrated in a remarkably sublinear manner (Figure 2). Interestingly, this phenomenon was recapitulated at the level of synaptic inputs in the voltage-clamp experiment, but to a lesser extent (Figure 3B-C). The polysynaptic response dependency of the sublinearity (Figure 3D-E) suggest that the inhibitory inputs are the causal mechanism. Because we clamped the membrane potential near the chloride equilibrium potential, where GABAergic transmission on the postsynaptic neurons does not affect the synaptic current, the observed sublinearity must be presynaptically targeted. At the same time, however, the inhibitory input showed typical features of feedforward inhibition on the postsynaptic neurons as the inhibitory current was temporally delayed and was excitatory input dependent (Supplementary Figure 4, 5). To support these findings, we identified that inhibitory neurons receiving the POm input modulated the vM1-driven excitatory inputs presynaptically and postsynaptically (Figure 4). Under physiological conditions, where inhibitions on both sides of the synapses are active, a greater degree of sublinearity can be expected, as demonstrated by the dendritic Ca^2+^ response.

Our immunohistochemical analysis suggested that the feedforward inhibition can be attributed to the PV and SST neurons. In particular, SST neurons are in a good position to exert mutual inhibition between POm and vM1 inputs on distal dendrites. Our anterograde trans-synaptic tracing results suggested overlapping yet distinct populations of SST neurons that received either POm or vM1 inputs (Figure 5). While PV neurons form synapses near the soma of the target neurons and inhibit the action potential generation of the entire neuron, a subset of the SST neurons, called the Martinotti cells, project axons to L1 and form extensive horizontal arborizations in L1 (Jiang et al., 2015; Karube et al., 2004; McGarry et al., 2010; Wang et al., 2004). The inhibitory effects of the SST neurons can be spatially constrained. Reportedly, inhibition by SST can often occur at the level of individual dendritic spines in pyramidal neurons (Chiu et al., 2013; Higley, 2014). In a recent electron microscopy-based connectomic study, Martinotti cells were shown to target the dendrites of neurons more often when they had different response properties (Kuan et al., 2022). Martinotti cells in the deep layer of vS1 are activated during whisking, although some SST neurons are involved in the canonical disinhibition circuit supporting gain regulation by context or attention (Fu et al., 2014; Gentet et al., 2012; Lee et al., 2013; Muñoz et al., 2017). Finally, unlike the activity of other types of inhibitory neurons, that of SST neurons is selectively tuned for specific sensory inputs (Ma et al., 2010) or behavior (Adler et al., 2019). Minimal trans-synaptic labeling in VIP neurons was unexpected (Figure 5) considering that previous studies have shown strong POm and vM1 inputs in VIP neurons (Audette et al., 2018; Lee et al., 2013; Sermet et al., 2019). We believe that the efficiency of trans-synaptic transmission might differ in a cell type-specific manner.

If POm inputs inhibit the vM1 inputs on distal dendrites and vice versa, how would the two inputs be integrated? We believe that the dendritic integration depends on the strength of the inputs. When the inputs are moderately strong and not strong enough to evoke firing in inhibitory neurons, the inputs depolarize the dendrites cooperatively (Schiller et al., 1997; Yuste et al., 1994). However, when one of the inputs is repeated at a high frequency or becomes strong enough to produce action potentials in inhibitory neurons, the feedforward inhibition is activated, which inhibits the other input. Although we leave the critical experiment examining the physiological relevance of the wiring to future work, we propose that such a flexible integration strategy could function as an electrical fuse. Current influx over the threshold for dendritic spikes is not only pointless but can also be hazardous to distal dendrites. Excessive increases in intracellular Ca^2+^ concentration generate free radical species and eventually lead to neuronal cell death in general (Duan et al., 2007). Notably, a degenerative cascade of cell death signaling originating from the distal dendrites has been reported upon the increased Ca^2+^ concentration in the dendrites (Shuttleworth and Connor, 2001). Furthermore, distal dendrites are susceptible to excessive synaptic inputs for the following reasons. Distal dendrites have high input impedance because of the small diameter of the dendritic segments (Branco and Häusser, 2011); thus, relatively large voltage changes are generated by the same amount of synaptic current compared to that in proximal dendrites (Fiala et al., 2007). Furthermore, when the two inputs are physically close to each other, such as the synapses from vM1 and POm (Kim et al., 2022), two temporally synchronized inputs can add extra voltage and cause Ca^2+^ changes by opening voltage-gated Ca^2+^ channels and activating NMDA receptors (Gasparini et al., 2004; Losonczy and Magee, 2006). Therefore, it is tempting to conclude that feedforward inhibition on the distal dendrites may function as a safety mechanism to maintain the inputs under a certain level. It is an interesting experimental model to address how the integration modes of multiple inputs may switch depending on the strengths of the inputs and how critical the function of SST neurons in the integration mode switching is. The current study has an explicit limitation—it does not provide direct evidence of how dendritic responses related to different behavioral parameters (i.e., whisking and object-touching) may interfere with each other. Instead, we examined how the inputs with various anatomical sources are integrated, assuming different brain regions are responsible for different information processing, and thus, inputs from different origins will convey different information. Nevertheless, we anticipate our results will broaden our understanding of the operation principles of the neocortex and help us comprehend how failure of the operation will lead to deficits in sensory processing in the diseased brain.

## Acknowledgments

This work was supported by the KBRI intramural research program (KBRI 22-BR-01-01, 22-BR-03-01, and 22-BR-04-04), DGIST R&D Program (21-IJRP-01), Korea Research Institute of Bioscience and Biotechnology(KRIBB) Research Initiative Program (KGM5282221, KGM4562222) and the National Research Foundation of the Ministry of Science and ICT (NRF-2022R1A2C1004216 and 2017M3A9G8084463).

## Supplementary figure legends

**Supplementary Figure 1.**
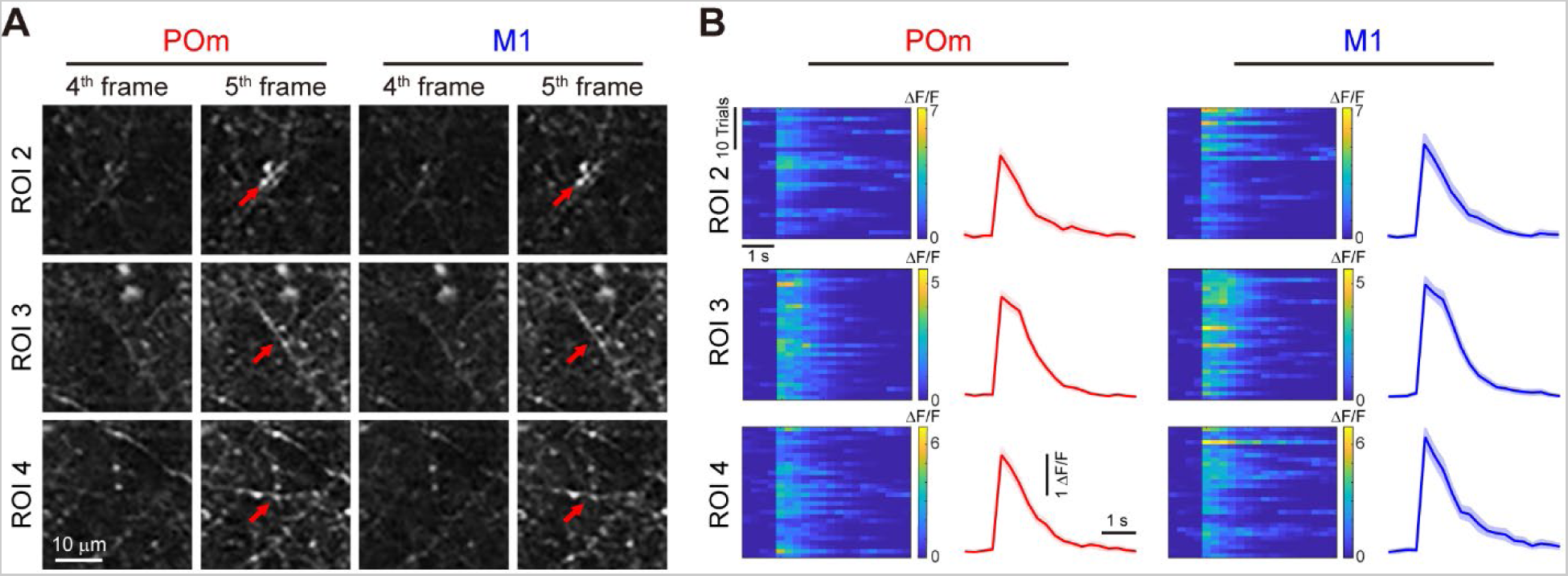
Further examples of the calcium deflection of ROIs. **A.** Example images of baseline (4^th^), immediately after stimulation (5^th^), and tail responses (10^th^ and 20^th^ frames) from a dendritic branch responding to both POm (left) and vM1 (right) stimulation. Images are from the red box 2–4 in C. **B**. Color-coded ΔF/F_0_ for repetitive 30 trials and averaged ΔF/F by the stimulation of POm and vM1. The shaded areas represent the standard errors of the responses.

**Supplementary Figure 2.**
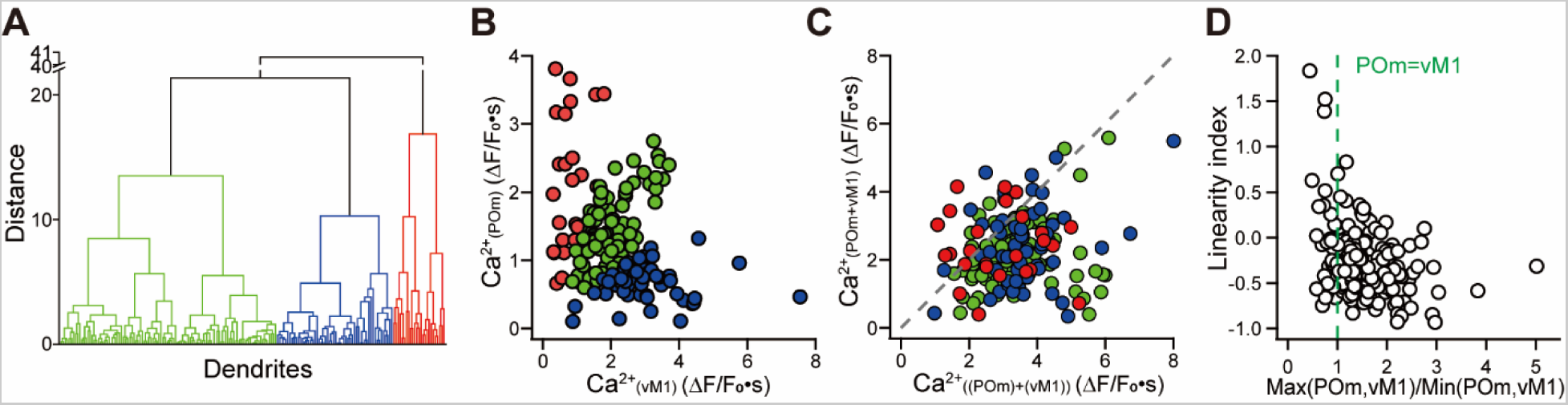
Linearity index as a function of input strength differences. **A.** Hierarchical clustering analysis of the relative strengths between Ca^2+^ responses by POm and those by vM1. Clustering was analyzed in a three-dimensional space with Ca^2+^ responses by POm(Ca^2+^_POm_), those by vM1 (Ca^2+^_vM1_), and the ratio of the two responses (atan(Ca^2+^_POm_/ Ca^2+^_vM1_) in degree) in each axis. **B.** Dendritic patches were divided into three distinctive groups on the Ca^2+^_POm_ and Ca^2+^_vM1_ plane. Depending on the relative strength of the inputs, we have plotted the branches in red (branches with strong Ca^2+^_POm_), green (2xCa^2+^_POm_ ≅ Ca^2+^_vM1_), and blue (branches with strong Ca^2+^_vM1_). **C.** Population Ca^2+^ deflection by simultaneous POm and vM1 stimulation as a function of the linear expected value by individual stimulations (n = 185) with the color codes demonstrating the relative strength of the Ca^2+^_POm_ and Ca^2+^_vM1_. **D.** Distribution of linearity index (Ca^2+^_POm+vM1_ – (Ca^2+^_POm_ + Ca^2+^_vM1_)}/ (Ca^2+^_POm_ + Ca^2+^_vM1_)) as a function of relative strength of the two inputs.

**Supplementary Figure 3.**
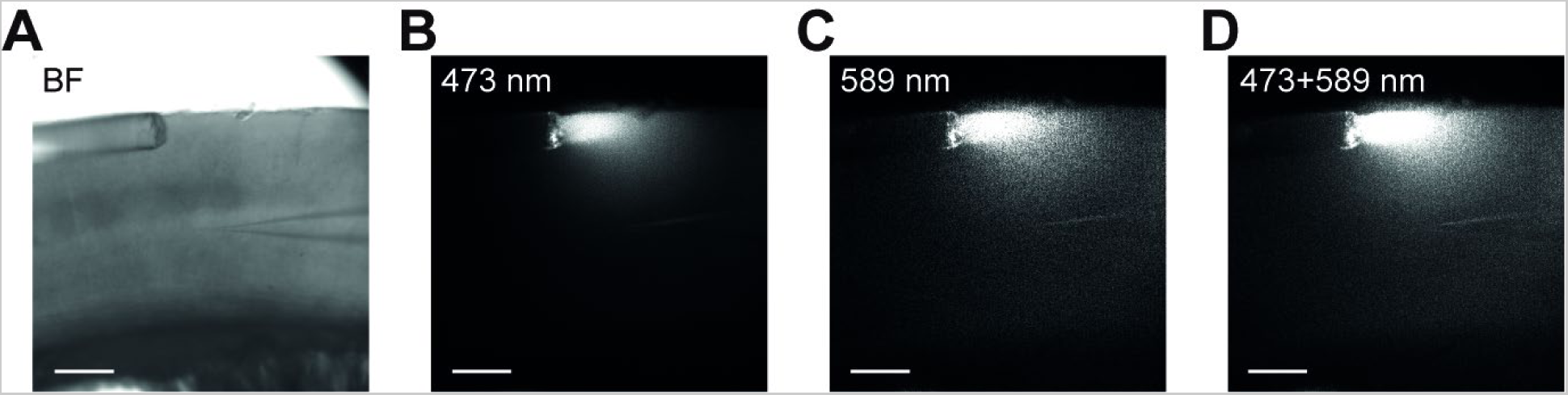
Photostimulation of distal inputs. **A.** Bright-field image of the experimental setup. **B-D.** The illuminated excitation light on the top layer of vS1. Note that light was focused on the distal dendrites of the recorded neuron.

**Supplementary Figure 4.**
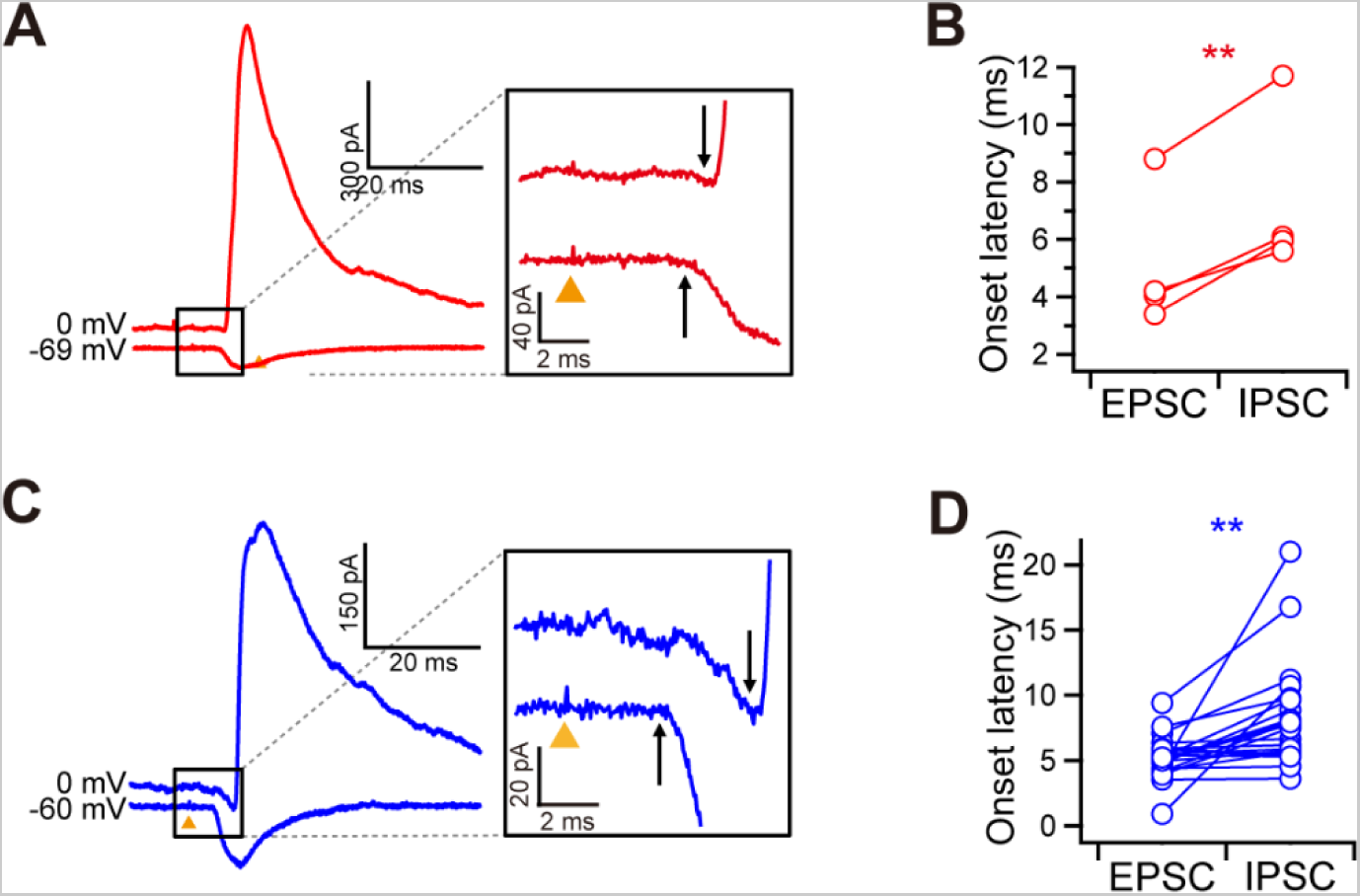
Feedforward inhibition in distal dendrites. **A and C.** Example traces of excitatory and inhibitory postsynaptic currents measured at 0 mV and the reversal potential of Cl^−^ evoked by optogenetic stimulation of POm (A) and vM1 (C). (Inset) Magnified at the onset of EPSCs and IPSCs. **B and D**. Onset latencies of excitatory and inhibitory synaptic currents by POm (B) and vM1 (D) stimulation.

**Supplementary Figure 5.**
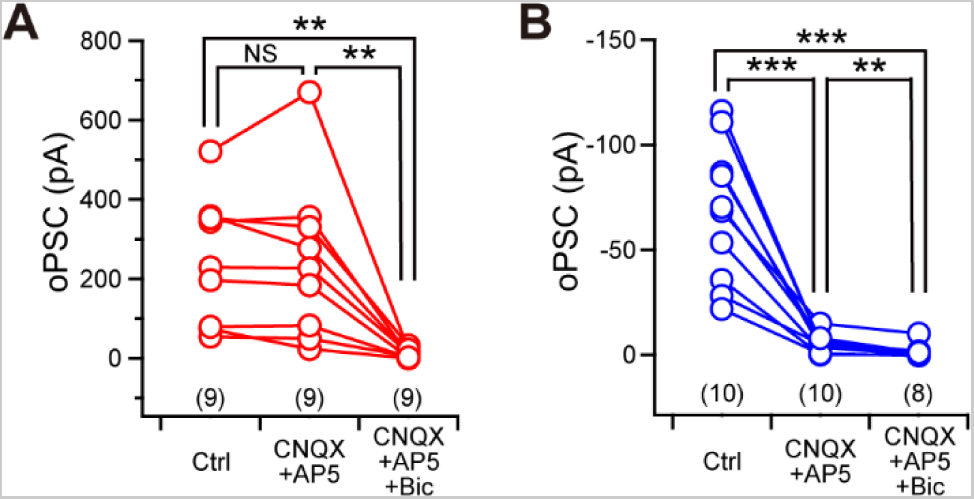
Voltage and receptor dependency of the optogenetically evoked postsynaptic current. **A.** oPSCs stimulated by POm-connected inhibitory neurons (POm-IN) measured at 0 mV and pharmacological sensitivity to glutamate receptor antagonists and GABA. Note that the amplitude of the oPSCs was not changed by the glutamate receptor antagonist CNQX (10 μM) but was completely abolished by the GABA_A_ antagonist bicuculline (20 μM). **B.** Pharmacological dependence of oPSC evoked by optogenetic activation of vM1 by chrimsonR measured at approximately −70 mV.

